# A widespread family of WYL-domain transcriptional regulators co-localises with diverse phage defence systems and islands

**DOI:** 10.1101/2021.12.19.473342

**Authors:** David M. Picton, Joshua D. Harling-Lee, Samuel J. Duffner, Sam C. Went, Richard D. Morgan, Jay C. D. Hinton, Tim R. Blower

## Abstract

Bacteria are under constant assault by bacteriophages and other mobile genetic elements. As a result, bacteria have evolved a multitude of systems that protect from attack. Genes encoding bacterial defence mechanisms can be clustered into “defence islands”, providing a potentially synergistic level of protection against a wider range of assailants. However, there is a comparative paucity of information on how expression of these defence systems is controlled. Here, we functionally characterise a transcriptional regulator, BrxR, encoded within a recently described phage defence island from a multidrug resistant plasmid of the emerging pathogen *Escherichia fergusonii*. Using a combination of reporters and electrophoretic mobility shift assays, we discovered that BrxR acts as a repressor. We present the structure of BrxR to 2.15 Å, the first structure of this family of transcription factors, and pinpoint a likely binding site for ligands within the WYL-domain. Bioinformatic analyses demonstrated that BrxR homologues are widespread amongst bacteria. About half (48%) of identified BrxR homologues were co-localised with a diverse array of known phage defence systems, either alone or clustered into defence islands. BrxR is a novel regulator that reveals a common mechanism for controlling the expression of the bacterial phage defence arsenal.

## Introduction

Bacteriophages outnumber bacterial prey by about ten-fold (1, 2). The estimated ≥10^30^ bacteriophages (phages) on Earth (1, 2) cause infections at a rate of 10^25^ per second (3). To contend with this extreme selection pressure, bacteria have evolved varied modes of defence against phages and other mobile genetic elements (4–6). Well-established examples of defence systems include restriction-modification (R-M) (7–10), abortive infection (11) and CRISPR-Cas (12) systems. R-M systems have been shown to cluster in “immigration control regions” (13). Recent comparative genomic analyses have demonstrated how diverse defence systems also commonly cluster into “defence islands” (14, 15). The “guilt-by-association” approach has allowed gene functions to be inferred from defence islands, and has identified novel defence systems (16). Coupled with renewed interest in technological spin-offs from these systems, and the rise of phage therapy to treat bacterial infections, multiple new systems have been identified, including Bacteriophage Exclusion (BREX) (17), CBASS (18), BstA (19), retrons (20), viperins (21) and pycsar (22). As multiple diverse systems have been assembled into a single locus, expression of the various genes must be meticulously regulated to reduce any impacts on host fitness whilst maximising the response to phages, and other mobile genetic elements.

It has been postulated that WYL-domain containing proteins act as ligand-binding regulators of phage defence system expression (23). WYL-domains (named after three conserved amino acids), are only found in prokaryotes and are part of the Sm/SH3 superfold family, which is itself subsumed by the larger “small β-barrel” family (24). Sm proteins are responsible for eukaryotic snRNP complexes and were first discovered as autoantigens in cases of lupus (using sera from a patient named Stephanie Smith) (25), whilst SH3 (Src-homology 3) domains are adaptor domains with diverse roles in eukaryotic cell signalling (26). In prokaryotes, the Sm homologue Hfq uses the Sm/SH3 fold to bind RNAs (27), whilst other WYL-domains bind proteins, peptides, DNA and oligosaccharides (23).

A handful of studies have begun to demonstrate that WYL-domain containing proteins regulate diverse processes in prokaryotes: Sll7009 from *Synechocystis* 6803 represses the CRISPR subtype I-D locus (28); DriD from *Caulobacter crescentus* activates expression of SOS-independent DNA damage response mediators (29); PIF1 helicase from *Thermotoga elfii* has a ssDNA-binding WYL-domain that couples ATPase activity to DNA unwinding (30); the RNA cleavage activity of a Type VI Cas13d protein from *Ruminococcus* is stimulated by a WYL-domain protein named WYL1 (31), which binds ssRNA with high affinity (32); and in Mycobacteria, PafBC is a transcriptional activator of DNA damage response genes (33). Most recently, WYL-domain proteins were found associated with phage defence islands within integrative conjugative elements of *Vibrio cholerae* (34).

We recently characterised a multi-functional phage defence island containing a BREX system (17) and the BrxU GmrSD-family (35) type IV restriction enzyme, encoded on a multidrug resistant plasmid of the emerging animal and human pathogen *Escherichia fergusonii* (36, 37). These two systems provide complementary protection against a wide range of environmental coliphages (37). This defence island encodes a WYL-domain containing protein, BrxR, which was hypothesised to act as a transcriptional regulator. Here, we present functional and structural characterisation that identifies BrxR as the first member of a large family of transcriptional regulators. BrxR homologues are widely associated with diverse phage defence systems and islands. Our findings suggest a possible common thematic approach for the regulation of phage defence systems that may involve a signalling molecule acting as a secondary messenger.

## Materials and Methods

### Bacterial strains and culture conditions

Total genomic DNA (gDNA) was obtained for *E. fergusonii* ATCC 35469 from ATCC. *E. coli* strains DH5α and BL21 (DE3) (ThermoFisher Scientific) were grown at 37 °C, either on agar plates or shaking at 220 rpm for liquid cultures. Luria broth (LB) was used as the standard growth media for liquid cultures, and was supplemented with 0.35% w/v or 1.5% w/v agar for semi-solid and solid agar plates, respectively. Growth was monitored using a spectrophotometer (WPA Biowave C08000) measuring optical density at 600 nm (OD_600_). When necessary, growth media was supplemented with ampicillin (Ap, 50 µg/ml), tetracycline (Tc, 10 µg/ml), isopropyl-β-D-thiogalactopyranoside (IPTG, 1 mM), L-arabinose (L-ara, 0.1% or 0.01% w/v), or D-glucose (D-glu, 0.2% w/v).

### Use of environmental coliphages

*E. coli* phages Pau, Trib and Baz were isolated previously from freshwater sources in Durham, UK (37). To make lysates, 10 μl of phage dilution was mixed with 200 μl of *E. coli* DH5α overnight culture and mixed with 4 ml of sterile semi-solid “top” LB agar (0.35% agar) in a sterile plastic bijou. Samples were poured onto solid LB agar plates (1.5% agar) and incubated overnight at 37 °C. Plates showing a confluent lawn of plaques were chosen for lysate preparations and the semi-solid agar layer was scraped off into 3 ml of phage buffer. 500 μl of chloroform was added and samples were vigorously vortexed and incubated for 30 min at 4 °C. Samples were centrifuged at 4000 x g for 20 min at 4 °C and the supernatant was carefully transferred to a sterile glass bijou. 500 μl of chloroform was added and lysates were kept at 4°C for long term storage.

### DNA isolation and manipulation

PCR amplicons and plasmids were purified using Monarch DNA kits (NEB). PCR, restriction digests, ligations, transformations and agarose gel electrophoresis were performed using standard molecular biology techniques. Constructed plasmids were confirmed via sequencing with an Abi 3370 DNA sequencer. The pSAT1-LIC-*brxR*^+^ expression construct adds a cleavable N-terminal His_6_-SUMO tag. Primers TRB878 and TRB879 were used to amplify *brxR* from pEFER (gene pEFER_0020) for insertion into pSAT1-LIC (38) to produce pTRB446 via Ligation Independent Cloning (LIC) (**Supplementary Table S1**). Primers TRB876 and TRB877 were used to amplify *brxR* from pEFER which was inserted into pBAD30 (39) to produce pBAD30-*his*_*6*_*-brxR* (**Supplementary Table S1**).

### Efficiency Of Plating assays

*E. coli* DH5α were transformed with pBAD30-*his*_*6*_*-brxR* and transformants were used to inoculate overnight cultures. Serial dilutions of phages Pau, Trib, and Baz (37) were produced ranging from 10^−3^ to 10^−10^. 200 µl of overnight culture and 10 µl of phage dilution were added to 3 ml top LB agar and plated on solid LB agar supplemented with 0.2% D-glu or 0.1% L-ara, to repress or induce *brxR* expression from pBAD30 constructs, respectively. Plates were incubated overnight before plaque forming units (pfu) were counted on each plate. Efficiency of plating (EOP) values were calculated by dividing the pfu of the L-ara-containing plates by the pfu of the D-glu-containing plates. Data are the mean and standard deviation of three independent replicates.

### β-galactosidase assays

Putative promoter regions (R1-12) were ligated into the promoterless *lacZ* fusion plasmid, pRW50 (40) (**Supplementary Table S1**). *E. coli* DH5α was then co-transformed with one of the *lacZ* reporter constructs (or pRW50 as a vector control) and either pBAD30 or pBAD30-*his*_*6*_*-brxR*. Transformants were used to inoculate overnight cultures, supplemented with 0.2% D-glu or 0.01% L-ara, to repress or induce *brxR* expression from pBAD30 constructs, respectively. These were then used to seed 80 µl microplate cultures at an OD_600_ of either 0.05 (for cultures containing D-glu) or 0.1 (for cultures containing L-ara). These cultures were then grown to mid-log phase in a SPECTROstar Nano (BMG Labtech) plate reader at 37°C with shaking at 500 rpm. Cultures were then supplemented with 120 µl master mix (60 mM Na_2_HPO_4_, 40 mM NaH_2_PO_4_, 10 mM KCl, 1 mM MgSO_4_, 36 mM β-mercaptoethanol, 0.1 mg/ml T7 lysozyme, 1.1 mg/ml ONPG, and 6.7% PopCulture Reagent (Merck Millipore)). Initial OD_600_ readings were taken, and OD_420_ and OD_550_ readings were then taken every minute for 30 min, at 37 °C with shaking at 500 rpm. Miller Units (mU) were generated as described (41). The plotted data are the normalised mean and standard deviation of three independent replicates.

### Protein expression and purification

*E. coli* BL21 (DE3) was transformed with pSAT1-LIC-*brxR* and a single colony was use to inoculate a 25 ml overnight culture of LB, supplemented with Ap and grown overnight. Overnight cultures were used to inoculate 12 L of 2x YT media in 2 L baffled flasks, each containing 1L of culture. Cultures were grown at 37 °C shaking at 180 rpm until an OD_600_ of ∼0.6, at which point cultures were supplemented to a concentration of 1 mM IPTG to induce expression. Cultures were incubated overnight at 16 °C and cells were pelleted at 4500 x g for 30 min at 4 °C. Cell pellets were resuspended in 50 ml of ice-cold A500 (20 mM Tris–HCl pH 7.9, 500 mM NaCl, 10 mM imidazole and 10% glycerol) and used immediately or flash frozen in liquid nitrogen and stored at -80 °C.

Pellets were lysed via sonication and centrifuged at 45000 x g at 4 °C for 30 min. All clarified cell lysates were passed over a 5 ml HisTrap HP column (Cytiva), and washed with 50 ml of A500. Bound BrxR was further washed with 50 ml of W500 (20 mM Tris–HCl pH 7.9, 500 mM NaCl, 40 mM imidazole and 10% glycerol) and eluted from the column in B500 (20 mM Tris–HCl pH 7.9, 500 mM NaCl, 250 mM imidazole and 10% glycerol). Imidazole was removed via dialysis back into A500 and the sample was treated with hSENP2 SUMO protease overnight at 4 °C to remove the fusion tag. The resulting untagged BrxR was loaded on to a second 5 ml HisTrap HP column and the flowthrough was collected and concentrated to 2 ml. BrxR was further purified via size exclusion through a Sephacryl S-300 HR gel filtration column in preparative SEC buffer (20 mM Tris–HCl pH 7.9, 500 mM KCl and 10% glycerol). Fractions were analysed via SDS-PAGE to assess content and purity, and peak fractions were pooled. BrxR was either dialysed into Xtal buffer (20 mM Tris–HCl pH 7.9, 200 mM NaCl and 2.5 mM DTT) for use in crystallisation, or was supplemented with glycerol to a final concentration of 30 % (w/v) for biochemical assays and stored at -80 °C following flash freezing in liquid nitrogen.

### Electrophoretic Mobility Shift Assays

An inverted repeat (IR) region was identified within the R7 promoter region upstream of *brxR*. Probes were synthesised using artificial templates (IDT) containing the target region and a 3′ common region corresponding to the start of *lacZ* within pRW50. Templates consisted of either the WT sequence, or mutant sequences which replaced one or both of the IRs with polycytosine (**Supplementary Table S1**). Incorporation of a common region permitted the use of a single fluorescein-tagged reverse primer TRB1068, in conjunction with the respective forward primer, to produce fluorescein-tagged WT and IR mutant probes via PCR. Probes were purified via gel extraction and quantified via Nanodrop. DNA-binding reactions were performed in 10 μl volumes, containing 1 μl of 2500 fmol of labelled probe, 1 μl of 1 μg/μl poly (dI-dC), 2 μl of 5x EMSA buffer (50 mM Tris–HCl pH 7.9, 750 mM KCl, 2.5 mM EDTA pH 8.0, 1 mM DTT, 0.5% Triton X-100, 65% glycerol), 1 μl of BrxR and made up to 10 μl with water. Specific competitor samples used a 20-fold excess of unlabelled probe, and non-specific competitor samples used a 20-fold excess of the *rv2827c* promoter region from *Mycobacterium tuberculosis* (41) (**Supplementary Table S1**).

Samples were incubated at 25 °C for 30 min before being loaded onto 7% native PAGE gels in 0.5x TBE (45 mM Tris-borate pH 8.0, 1 mM EDTA). Gels were pre-ran at 150 V for 120 min for 2 hrs at 4 °C. Gels were imaged using an Amersham Bioscience Typhoon 9400 in fluorescence mode, emission filter 526 SP. Band intensities of the unbound probe were enumerated using ImageJ. Fractional saturation corresponding to the amount of unbound probe, Y, was calculated using Y = 1-(I_T_/I_C_), where I_T_ is the band intensity of the unbound probe in test lanes and I_C_ is the band intensity probe in the control lane at 0 mM BrxR. Dissociation coefficients (*K*_d_) were calculated from saturation plots using non-linear regression. Data shown are mean values from triplicate experiments and are plotted with standard error of mean.

### Analytical gel filtration

A Superdex 200 Increase (S200i) GL 5/150 (Cytiva) was connected to an ÄKTA Pure system (Cytiva) and equilibrated by running through 2 column volumes of filtered MQ water and 5 column volumes of analytical SEC buffer (20 mM Tris–HCl pH 7.9 and 150 mM NaCl) at 0.175 ml/min. It was then calibrated using standard calibration kits (Cytiva). Calibration curves were used to calculate the oligomeric state of BrxR according to its elution volume. A 50 μl sample containing 5 μl of 1000 nM BrxR, 5 μl of 5x FPLC sample buffer (100 mM Tris-HCl pH 7.9, 750 mM KCL, 20 % (w/v) glycerol) and made up with water. Samples were loaded onto to the S200i via Hamiliton syringe into a 10 μl loop. Samples were injected onto the S200i and two column volumes of analytical SEC buffer were used to elute BrxR at a flow rate of 0.175 ml/min.

### Protein crystallisation and structure determination

BrxR was concentrated to 10 mg/ml and crystallisation trials were set using a Mosquito Xtal3 robot (Labtech) with commercial screens (Molecular Dimensions). Drops were set at both 1 : 1 and 2 : 1 (protein : precipitant) ratios at 18 °C and crystals appeared overnight in Pact Premier F8 (0.2 M Na_2_SO_4_, 0.1 M Bis-Tris propane pH 7.5 and 20 % w/v PEG 3350). Crystals were reproduced manually in 2 μl drops and harvested in nylon cryoloops. Crystals were soaked in Cryo solution (20 mM Tris-HCl pH 7.9, 150 mM NaCl, 2.5 mM DTT and 80% glycerol) and stored in liquid nitrogen. Diffraction data were collected on I04 at Diamond Light Source (DLS). Four datasets collected at 0.9795 Å were merged to produce a single dataset using the DIALS pipeline in iSpyB (DLS). Data scaling was performed using AIMLESS (42). Phases were obtained by molecular replacement using an AlphaFold (43) model of BrxR in PHASER (PHENIX) (44), to produce a starting model which was then further built using BUCCANEER (45). Iterative refinement was performed using PHENIX and manually edited in COOT (46). Structure quality was assessed using PHENIX, COOT and the wwPDB validation server, and BrxR was solved to 2.15 Å. Structural figures were produced in Pymol (Schrödinger).

### Comparative Genomic Analyses

The protein sequences and features of 3,828 reference and representative prokaryote sequences of “complete” or “chromosome” quality were downloaded from RefSeq using ncbi-genome-download v0.2.9 (https://github.com/kblin/ncbi-genome-download), in September 2021. A BLAST database of 13,499,153 proteins was constructed, and the protein sequence of BrxR (pEFER_0020) queried against the database using blastp with default settings. All homologues with E-value < 1e-5 were identified. Marker genes were queried against the same database, using blastp at default settings, and all homologues with E-value < 1e-5 identified. A second query with a less stringent threshold of E-value < 1e-3 was also carried out. The location, proximity, and orientation to identified BrxR homologues was then determined (in-house R script). All BLAST analyses were carried out on a CLIMB server (47).

A representative phylogeny of the 3,828 genomes was downloaded from the NCBI Common Taxonomy Tree resource. The ete3 toolkit (48) provided taxonomic information for each genome. Trees were visualised in R using the ggtree package (49). UpSet plots were produced using the UpSetR R package (50).

## Results

### The pEFER phage defence island is regulated by BrxR

We previously characterised a phage defence island from *E. fergusonii* ATCC 35469 (37). The island is carried by pEFER, a multidrug-resistant, 55.15 kb, plasmid. By sub-cloning the 18 kb defence island, we demonstrated that the island provides phage defence via the complementary BREX system (17) and a GmrSD-family type IV restriction enzyme (35), named BrxU (37). We predicted that the third open reading frame of the ten-gene island was encoded a helix-turn-helix (HTH) domain, using PHYRE 2.0 (51). We hypothesised that this protein bound DNA to act as a transcriptional regulator of the defence island, and named it BrxR (**Figure 1A**). Bioinformatic analyses predicted a promoter upstream of *brxR* (52). Part of the transcriptional control of other BREX systems is mediated by promoters upstream of *brxA* and *pglZ* (17). We hypothesised that an additional promoter would lie upstream of *brxS* and *brxT*, to permit independent expression of the accessory genes we found to be necessary for BREX-mediated host genome methylation (37) (**Figure 1B**).

**Figure 1.**
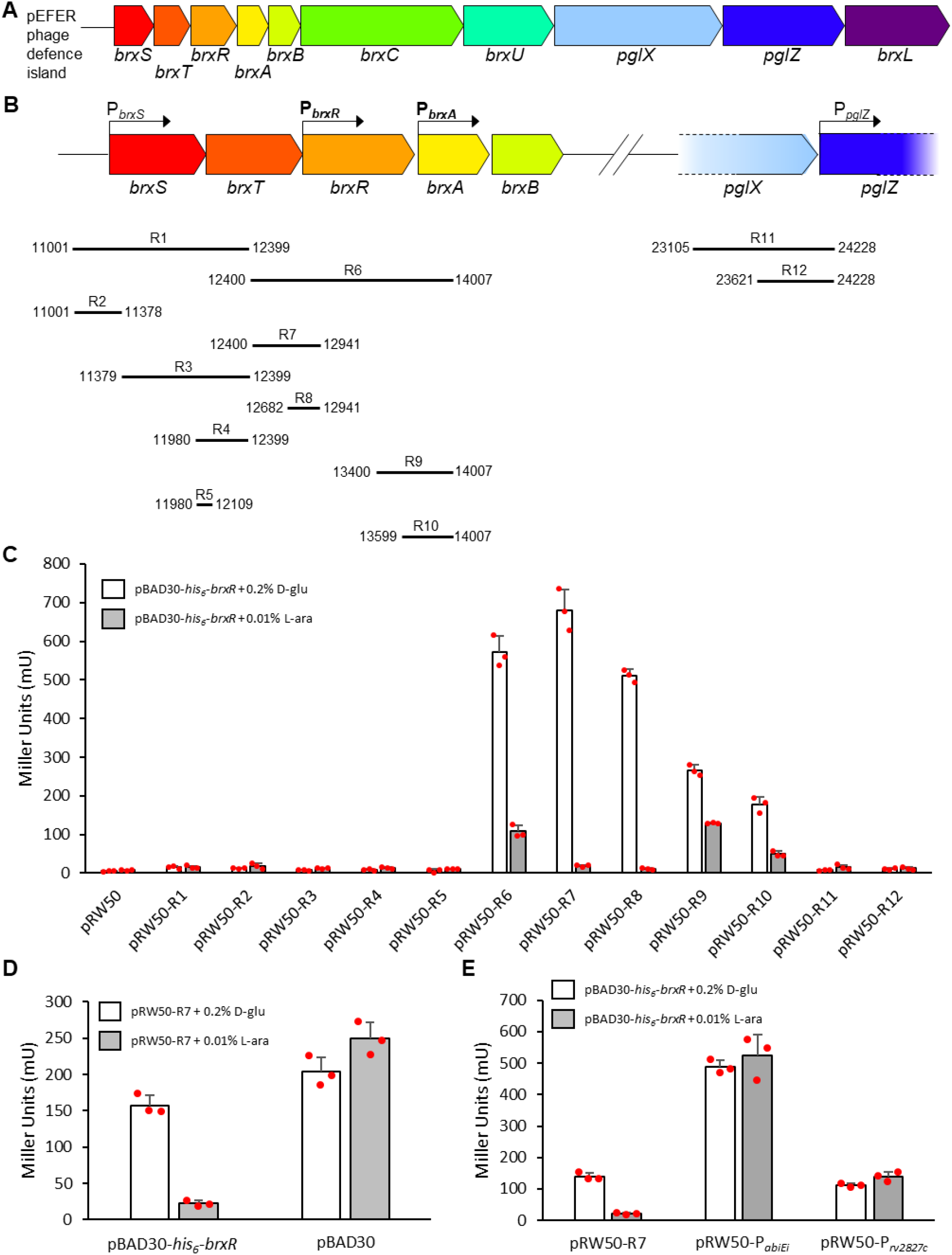
The pEFER phage defence island is regulated by BrxR at the transcriptional level. (**A**) Linear representation of the phage defence island of pEFER. (**B**) Transcriptional organisation of the pEFER phage defence island, showing putative promoters P_*brxS*_, P_*brxR*_, P_*brxA*_, and P_*pglZ*_, with an accurate alignment of experimental test regions R1-12 that were cloned into the promoterless *lacZ*-reporter plasmid, pRW50. (**C**) LacZ-reporter assays using constructs pRW50-R1-12 with and without the induction of BrxR from pBAD30-*his*_*6*_*-brxR*, showing activity from P_*brxR*_ and P_*brxA*_ (in bold within (**B**)), and repression by BrxR. (**D**) LacZ-reporter assays using pRW50-R7 with and without induction of pBAD30-*his*_*6*_*-brxR* or a pBAD30 vector control. (**E**) LacZ-reporter assays using active pRW50 promoter constructs with and without induction of BrxR from pBAD30-*his*_*6*_*-brxR*. Data are shown in triplicate, and error bars represent standard deviation of the mean.

To investigate the function of the hypothetical promoters, and to determine the ability of BrxR to regulate gene expression, regions of the pEFER defence island denoted R1-12 were cloned into pRW50 (40), which encodes a promoterless *lacZ* reporter gene (**Figure 1B**). Gene *brxR* was also cloned into pBAD30 (39) to permit L-arabinose-inducible expression of BrxR, yielding pBAD30-*brxR. E. coli* DH5α was co-transformed with either pRW50 vector control or reporter plasmids, and pBAD30-*brxR*. These dual plasmid-carrying strains containing both a pRW50 reporter and pBAD30-*brxR* were grown either in the presence of D-glucose (D-glu), to repress *brxR* expression, or L-arabinose (L-ara), to induce *brxR* expression, and the resulting levels of β-galactosidase activity were determined (**Figure 1C**).

Of the four putative promoter regions, strong expression was observed from a promoter upstream of *brxR*, (P_*brxR*_), with weaker expression being observed from upstream of *brxA* (P_*brxA*_). Neither regions upstream of *brxS* nor *pglZ* showed measurable levels of transcriptional activity (**Figure 1C**). The induction of *brxR* reduced the expression from P_*brxR*_ and P_*brxA*_ (**Figure 1C**). Using pRW50-R7, we then confirmed that repression was due to expression of BrxR when compared to an empty pBAD30 vector control (**Figure 1D**). Finally, to confirm that BrxR-mediated repression of transcription was specific to the tested DNA regions, rather than reflecting a global activity of BrxR, we tested whether BrxR could repress expression from pRW50-based reporter plasmids carrying other promoters (41) (**Figure 1E**). BrxR-mediated repression only occurred for the pEFER-derived promoter carried by pRW50-R7 (**Figure 1E**). Collectively, these data indicate that BrxR is a transcriptional regulator of the pEFER phage defence island that negatively regulates expression.

We additionally tested whether the pBAD30-*brxR* plasmid provided any protection from phages that were previously shown to be susceptible to the pEFER defence island (37) (**Supplementary Figure S1**). BrxR alone had no impact on the ability of the tested phages to form plaques confirming BrxR to be a regulator of, but not a participant within, phage defence (**Supplementary Figure S1**). We have previously sub-cloned the defence island from pEFER, generating plasmid pBrxXL that demonstrated complementary defence through BREX and BrxU (37). Next, we aimed to test the impact of ablating *brxR* expression in the context of the pBrxXL plasmid. However, the putative *brxR* knockout transformants obtained by either golden gate assembly (53), or Gibson assembly (54), all contained extensive mutations in other parts of the defence island. Furthermore, when commissioned, the *brxR* knockout plasmid could not be generated commercially. Collectively, our findings imply that the repression provided by BrxR in the context of the pEFER defence island may both regulate phage defence and limit inherent toxicity associated with uncontrolled expression of the island.

### BrxR binds inverted DNA repeats

Our previously studied HTH transcriptional regulators were shown to bind inverted DNA repeats (41, 55). We examined regions R7 and R9 (**Figure 1B**) and found an 11 bp imperfect inverted repeat (containing a single base mismatch at the second position), with a 5 bp spacer between repeats, located between P_*brxR*_ and the ribosome binding site of *brxR* (**Figure 2A**). Curiously, no inverted repeats were identified upstream of P_*brxA*_.

**Figure 2.**
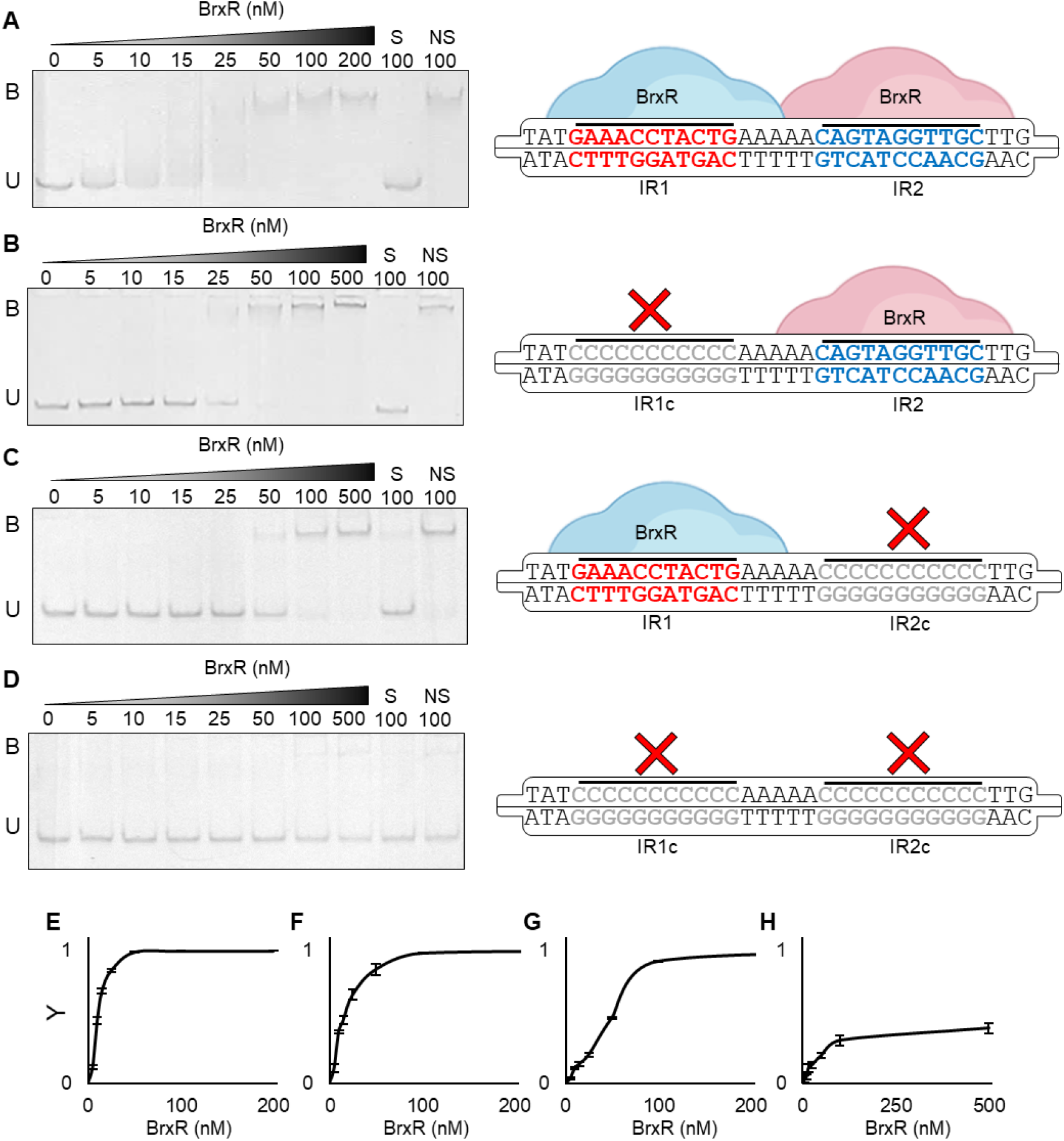
BrxR binds DNA inverted repeats *in vitro*. (**A-D**), Electrophoretic mobility shift assays (EMSAs) of titrated BrxR protein with dsDNA probes spanning pEFER nucleotide locations 12801-12870. Target probes were amplified to incorporate fluorescein and contain either the native promoter region (**A**) or substituted regions where either IR1 (**B**), IR2 (**C**) or both IRs (**D**) were replaced by polycytosine residues. Protein concentration is shown above each lane together with binding events (B -bound, U – unbound). Control lanes correspond to samples prepared with a 20-fold excess of unlabelled specific DNA (S) or non-specific DNA (NS), respectively. Experiments were run in triplicate and a representative gel from each experiment is shown. Each EMSA is accompanied with a schematic of the binding capacity of BrxR relative to the presence/absence of its target motif. IR mutations are shown in grey. Probe sequence diagrams are truncated to show only the IR regions of dsDNA probes. (**E-H**), Saturation curves were plotted using EMSA band intensity of unbound probe to determine Y values. Y values were calculated using Y = 1-(I_T_/I_C_), where I_T_ is the band intensity of the unbound probe in test lanes, and I_C_ is the band intensity probe in the control lane at 0 nM BrxR. Points plotted are mean values from triplicate data and error bars correspond to standard error of the mean. (**E**) Native promoter (**F-H**), Mutated promoter regions with polycytosine substitution of IR1 (**F**), IR2 (**G**) or both (**H**).

We tested the ability of BrxR to bind the repeats downstream of P_*brxR*_ by electrophoretic mobility shift assay (EMSA; **Figure 2A**). BrxR bound a labelled DNA probe that contained both inverted repeat 1 (IR1) and inverted repeat 2 (IR2), in a concentration-dependent manner (**Figure 2A**). The specific, S, control that contained a 20-fold excess of unlabelled probe, and the non-specific, NS, control, that contained a 20-fold excess of unlabelled probe from an unrelated *Mycobacterium tuberculosis rv2827c*-derived promoter (41), confirmed that the BrxR-DNA interaction was DNA-sequence specific (**Figure 2A**). BrxR-DNA binding generated a single shift of the labelled probe (**Figure 2A**), implying a single binding event. The presence of the two inverted repeats suggests that BrxR likely forms a stable dimer in solution that binds both IR1 and IR2 simultaneously. Replacing IR1 with a polyC tract still yielded a single binding event to IR2, albeit requiring greater concentrations of BrxR (**Figure 2B**). Similarly, replacing IR2 with a polyC tract had the same effect (**Figures 2C**). Replacing both IR1 and IR2 with polyC tracts prevented BrxR binding, unless at such high concentrations to allow non-specific DNA interactions (**Figures 2D**).

Quantification of complex formation in comparison to unbound probe generated *K*_d_ values for each binding event (**Figures 2E-H**). BrxR bound to the WT probe most tightly (*K*_d_ of 13.0 nM), followed by IR1c-IR2 (*K*_d_ of 24.0 nM), then IR1-IR2c (*K*_d_ 85.5 nM) (**Figures 2E-H**), suggesting that the mismatched base in IR1 reduced the affinity of BrxR binding. These data demonstrate that BrxR is a transcriptional regulator that binds inverted repeats to negatively regulate expression of phage defence genes.

### BrxR is the first example of a family of multi-domain dimeric transcriptional regulators

BrxR was expressed and purified as described (Materials and Methods), and its size in solution was determined by analytical size exclusion chromatography (analytical SEC) (**Figure 3A**). The elution volume indicated that BrxR forms a dimer in solution (**Figure 3A**), which was consistent with the observation of single binding events by EMSA, independent of there being one or two IRs (**Figure 2**).

**Figure 3.**
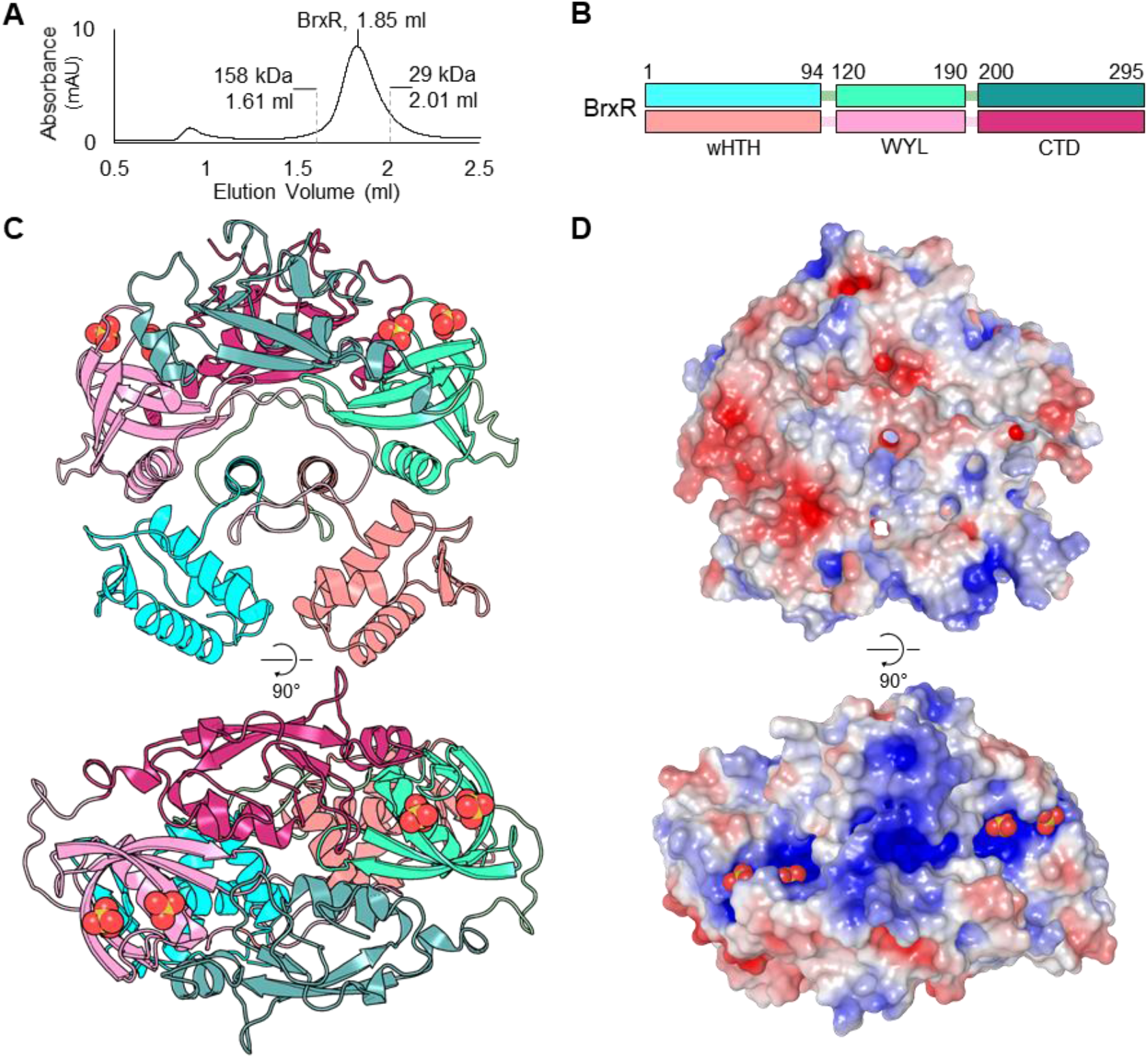
BrxR forms a dimer and exhibits significant surface electropositivity. (**A**) Size exclusion chromatography of BrxR resolved via a Superdex 200 increase GL 5/150 gel filtration column. BrxR elutes at an elution volume of 1.85 ml, corresponding to a mass twice its M_r_, indicating dimer formation. No additional peak is observed for residual monomers. Calibration standards are indicated. Organisation of the 3 domains within BrxR, separated by linker regions. Each protomer is coloured in either shades of cyan or pink, with domains indicated by the amino acid residue numbers shown. Cartoon overview of the BrxR dimer coloured corresponding to (**A**), presented in orthogonal views. Sulphate ions are represented as yellow and red spheres. (**D**) Electrostatic representation of surface BrxR charges in orthogonal views. The blue electropositive patches around the helices of the wHTH domains, and within the WYL domain surrounding the bound sulphate ions, are shown.

We solved the structure of BrxR by X-ray crystallography, to 2.15 Å (**Figure 3, Table 1**). The asymmetric unit contained four BrxR dimers, supporting our previous data. Each BrxR protomer consists of three domains (**Figure 3B**). BrxR comprises an N-terminal winged-HTH domain (residues 1-94) (56), followed by a WYL-domain (so called due to a previous analysis of conserved amino acids), which has been implicated as a potential ligand-binding domain with a role in phage defence (residues 120-190) (23), and a C-terminal dimerisation domain (residues 200-295) (**Figure 3B**).

**Table 1.** Crystallographic data collection and refinement statistics.

|  | BrxR Native |
| --- | --- |
| PDB ID Code | 7QFZ |
| Number of crystals | 1 |
| Beamline | Diamond I04 |
| Wavelength, Å | 0.9795 |
| Resolution range, Å | 82.35–2.15 (2.15–2.19) |
| Space group | P 4<sub>1</sub> 22 |
| Unit cell |  |
| *a b c* (Å) | 131.13 131.13 358.369 |
| *α β γ* (°) | 90.000 90.000 90.000 |
| Total reflections | 324782 |
| Unique reflections | 169928 |
| Multiplicity | 1.9 |
| Completeness (%) | 100 |
| Mean *I*/sigma(*I*) | 9.5 |
| *R*<sub>merge</sub> | 0.04 |
| *R*<sub>meas</sub> | 0.057 |
| CC<sub>1/2</sub> | 0.998 |
| *R*<sub>work</sub> | 0.216 |
| *R*<sub>free</sub> | 0.241 |
| No. of non-hydrogen |  |
| atoms | 19401 |
| Macromolecules | 80 |
| Ligands | 16 |
| Solvent | 884 |
| Protein Residues | 2276 |
| RMSD (bonds, Å) | 0.008 |
| RMSD (angles, °) | 1.16 |
| Ramachandran favoured (%) | 96.85 |
| Ramachandran allowed (%) | 3.15 |
| Ramachandran outliers (%) | 0 |
| Average B-factor | 59 |
| Macromolecules | 60.05 |
| Ligands | 63.06 |
| Solvent | 53.13 |

The wHTH domains are spaced ∼25 Å apart in the BrxR dimer, indicating additional movement is required to optimise the positions for interaction with the major grooves of target DNA (**Figure 3C**). The wHTH domains are exchanged between the protomers, such that upon exiting the wHTH the protein fold crosses to the other side of the dimer, to the WYL-domain. This cross-over begins with an α-helix aligned in parallel with that of the opposing protomer (residues 84-94), before entering a long loop section (residues 95-119) that circles round either end of both the central helices, interacting with all three domains of the opposing protomer around the circumference, before forming the WYL-domain. The first α-helix of each WYL also lines-up in parallel either side of the two central cross-over helices, to form a row of four parallel helices, alternating between protomers. The WYL-domains do not appear to directly interact, but the C-terminal dimerisation domains extend across like two left hands shaking, interacting through the opposing C-terminal domain through the palms, and with the opposing WYL-domain through an α-helix at the utmost tip of the protomeric dimerisation domain (residues 202-232) (**Figure 3C**). Two sulphate molecules are bound within each WYL-domain (**Figure 3C**). As the crystals were formed in conditions containing 0.2 M sodium sulphate, it is expected that the abundance of sulphate in the crystallisation condition allowed these ions to be resolved in the structure. Nevertheless, the position of the two sulphates corresponds to a solvent-exposed basic patch formed by each BrxR protomer (**Figure 3D**).

Protein sequences homologous to BrxR were selected with Consurf (57), and used for multiple sequence alignment and subsequent calculation of residue conservation. The conservation output was then mapped onto the BxrR surface (**Supplementary Figure S2A**). Interestingly, conservation showed a similar distribution to the electrostatic potential (**Figure 3D**), with greatest conservation in the DNA-contacting helices of the wHTH domain, the sulphate-binding residues of the WYL-domain, the central line of interfacing α-helices, and the protomer interface residues of the C-terminal dimerisation domain (**Supplementary Figure S3A**). The DALI server (58) was used to search the PDB for structural homologues of BrxR (**Supplementary Table S2**). The highest scoring hit reached a Z-score of only 11.3, indicating that there was no clear match to BrxR within the PDB. Of the obtained hits, each was shown to overlay either with the wHTH domain, a common DNA-binding motif (56), or the WYL-domain itself (33). No hits scoring above a Z-score of 4.0 matched the BrxR C-terminal dimerisation domain. We conclude that BrxR is the first solved structure of a new family of WYL-domain containing transcriptional regulators.

One other structure of a BrxR homologue that appears to regulate a phage defence island from *Acinetobacter* sp. NEB394 (BrxR_*Acin*_) has been solved by Luyten et al., in a study co-submitted with this article (59). We exchanged BrxR homologue structural models for comparison. A sequence-independent superposition of the two structures generated a Root Square Mean Deviation (RMSD) of 3.37-3.63 Å, depending on which of our modelled dimers of BrxR from *E. fergusonii* (from now on referred to as BrxR_*Efer*_) was used (**Supplementary Figure S2B**). Although the relatively low RMSD value suggests poor overall structural homology, both homologues have a similar arrangement of the same three domains, with variations in the relative positioning of each domain and secondary structure elements (**Supplementary Figure S2B**). For instance, whilst the wHTH domains remain ∼25 Å apart in the BrxR_*Acin*_ structure, they are tilted further out along the short axis of the dimer compared to BrxR_*Efer*_. Furthermore, the central parallel helices within BrxR_*Efer*_ are tilted in BrxR_*Acin*_, and the loop extending around and towards the protomeric WYL-domain of BrxR_*Acin*_ donates a β-strand to form an extended β-sheet with the opposing WYL domain as it passes.

Luyten et al. (59) also obtained a structure of BrxR_*Acin*_ in complex with DNA, having identified a similar set of target DNA inverted repeats, each of 10 bp (single mismatch at position 5) and separated by 5 bp. This DNA-bound structure shows that BrxR_*Acin*_ bends the target DNA, which allows interactions with the major grooves despite the spacing of only ∼25 Å between wHTH domains. When superposed against the BrxR_*Acin*_-DNA structure, the wHTH domains of BrxR_*Efer*_ also fit into the major grooves (**Supplementary Figure S2C**), but the winged β-sheet clearly clashes with the DNA backbone (**Supplementary Figure S2C, *inset***). This suggests that upon BrxR_*Efer*_ binding DNA, a conformational shift will be needed.

### WYL-domain of BrxR as a potential ligand sensor

WYL-domains have been proposed as ligand-binding domains that could act as sensors of phage infection to regulate phage defence systems (23). The fold of the WYL-domain from BrxR_*Efer*_ corresponds exactly with the expected features of the superfold Sm/SH3 family, itself a subset of the larger and pervasive small β-barrel (SBB) protein domain urfold family (24). The BrxR_*Efer*_ WYL-domain folds as an N-terminal α-helix, followed by five β-sheets (**Figure 4A**). The RT loop links sheets β1-β2 (numbered within this domain, not across the entire BrxR_*Efer*_ protein), the n-Src loop links sheets β2- β3, the distal loop links sheets β3-β4, and the short 3_10_ helix links sheets β4-β5 (**Figure 4A**). The top DALI hit (**Supplementary Table S2**) was the WYL-domain from PafBC (PDB 6SJ9) (33). The WYL-domains of BrxR_*Efer*_ and PafBC superpose with an RMSD of 0.662 Å, which is an overall good match, but there are distinct movements in the RT loop used by BrxR_*Efer*_ to bind sulphates (**Figure 4B**).

**Figure 4.**
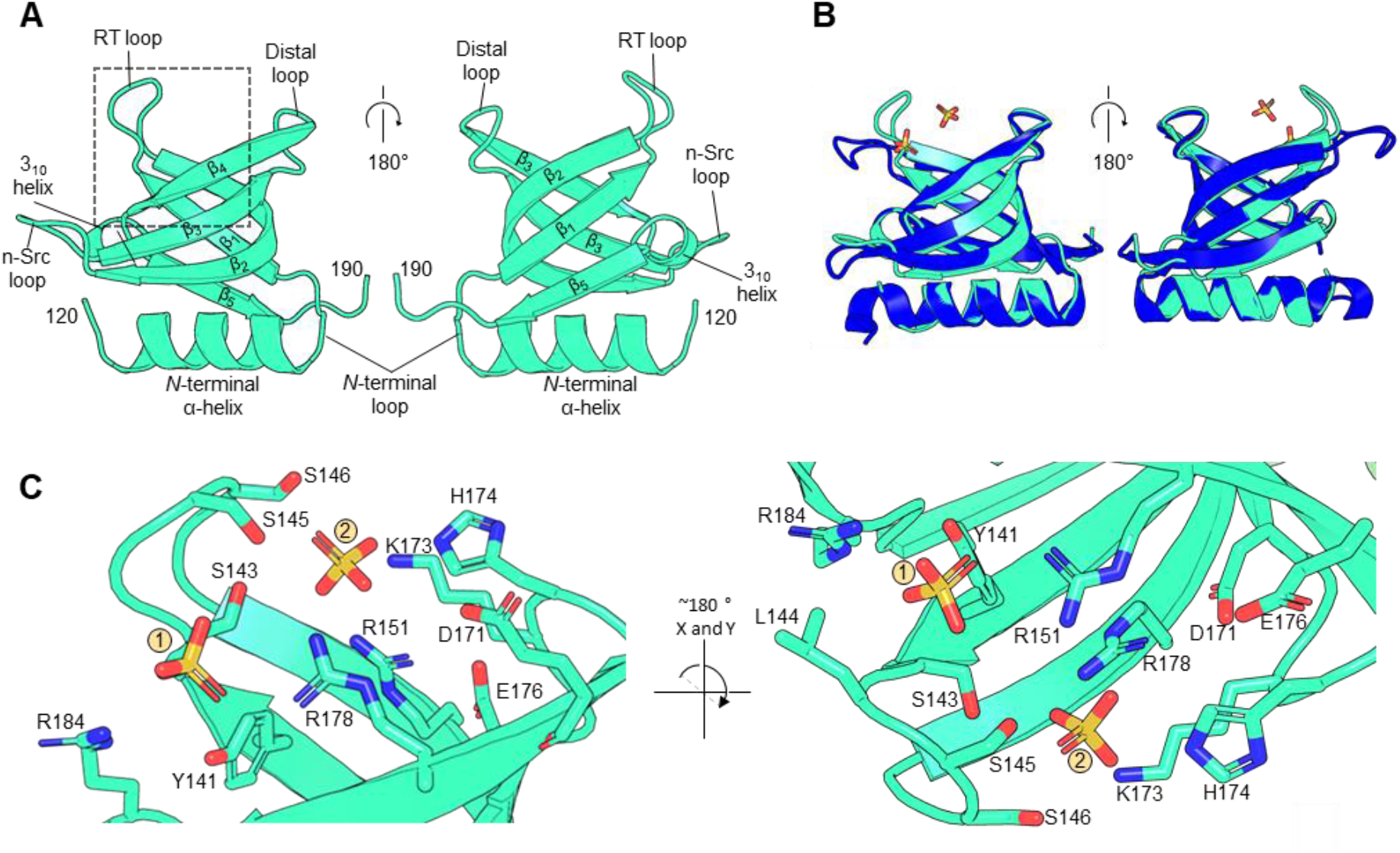
The BrxR WYL-domain shows ligand binding potential via extensive sidechain coordination. Orthogonal views are shown for each panel as indicated. (**A**) Close-up of the WYL domain of BrxR. Terminal residues are numbered, and secondary structural elements and loops for this domain are labelled. (**B**) Structural superposition of BrxR_*Efer*_ with the WYL-domain of PafBC (RMSD 0.662 Å; PDB 6SJ9) shows clear structural similarity. Differences are observed at the RT loop of BrxR_Efer_, which has moved inwards to bind the two sulphate ions. BrxR_*Efer*_ is shown in cyan and PafBC is shown in blue. (**C**) A close-up view of the dashed boxed area of (**A**) shows the hydrogen bond coordination of two sulphate ions bound within the WYL domain of BrxR_*Efer*_. Interacting sidechains extend from the core β-strands and intervening loops. Nitrogen atoms are shown in blue and oxygen atoms in red. Sulphate ions are shown as yellow (sulphur) and red (oxygen) sticks.

We noted that the overall arrangement of domains differs hugely between PafBC and BrxR_*Efer*_, and the C-terminal domains do not overlay, so they are from different protein families. Sm/SH3 domains are known to bind diverse polymeric ligands such as DNA, RNA, oligosaccharides, proteins and peptides (24). Detailed examination of the sulphate binding site in BrxR_*Efer*_ shows an abundance of polar groups that have captured the sulphate ions and could theoretically recognise other small molecule ligands (**Figure 4C**). These residues are found within the core β-strands but also the conserved loops, such as S143, S145 and S146 on the RT loop; K173 and H174 on the distal loop; and R184 on the 3_10_ α-helix (**Figure 4C**). We propose that ligand-binding at the WYL-domain basic patch could induce conformational changes to BrxR_*Efer*_ that relieve transcriptional repression.

### BrxR-family homologues are predominantly found in Proteobacteria

We wanted to investigate the extent of this newly identified BrxR family. Homologues were identified through bioinformatic searches of a protein database constructed from representative RefSeq genomes (see Materials and Methods), using BrxR_*Efer*_ as a search sequence with a conservative threshold of E-value < 1e-5. This threshold was chosen to exclude false positives associated with the prevalence of both wHTH and WYL-domains, the numbers of regulatory proteins in general, and the relative size of BrxR_*Efer*_. Our search identified 347 homologues within 281 genomes, including 59 genomes (57 proteobacteria, 1 firmicute, 1 planctomycete) encoding more than one BrxR-family protein. This corresponds to BrxR-family homologues in 7.79% of the 3,828 genomes in our representative dataset. All homologues were found in bacterial genomes, with no homologues identified in the 222 archaeal genomes.

We then considered the taxonomic distribution of the BrxR-family, and noted that 338/347 BrxR homologues were found throughout Proteobacteria (97.41% of total homologues), most commonly in *Pseudomonas* (24/347; 6.92%), *Shewanella* (18/347; 5.19%) and *Vibrio* (15/347; 4.32%) (**Figure 5**). Though widespread, no homologues were found within Deltaproteobacteria (**Figure 5**). BrxR homologues were found in 271 of 1589 proteobacter genomes within the dataset (**Figure 5**). Hits were additionally found in firmicutes (6 homologues in 607 genomes), spirochaetes (1 homologue in 49 genomes), planctomycetes (1 homologue in 62 genomes) and verrucomicrobia (1 homologue in 110 genomes). Collectively, these data show that the BrxR-family is widespread amongst Proteobacteria.

**Figure 5.**
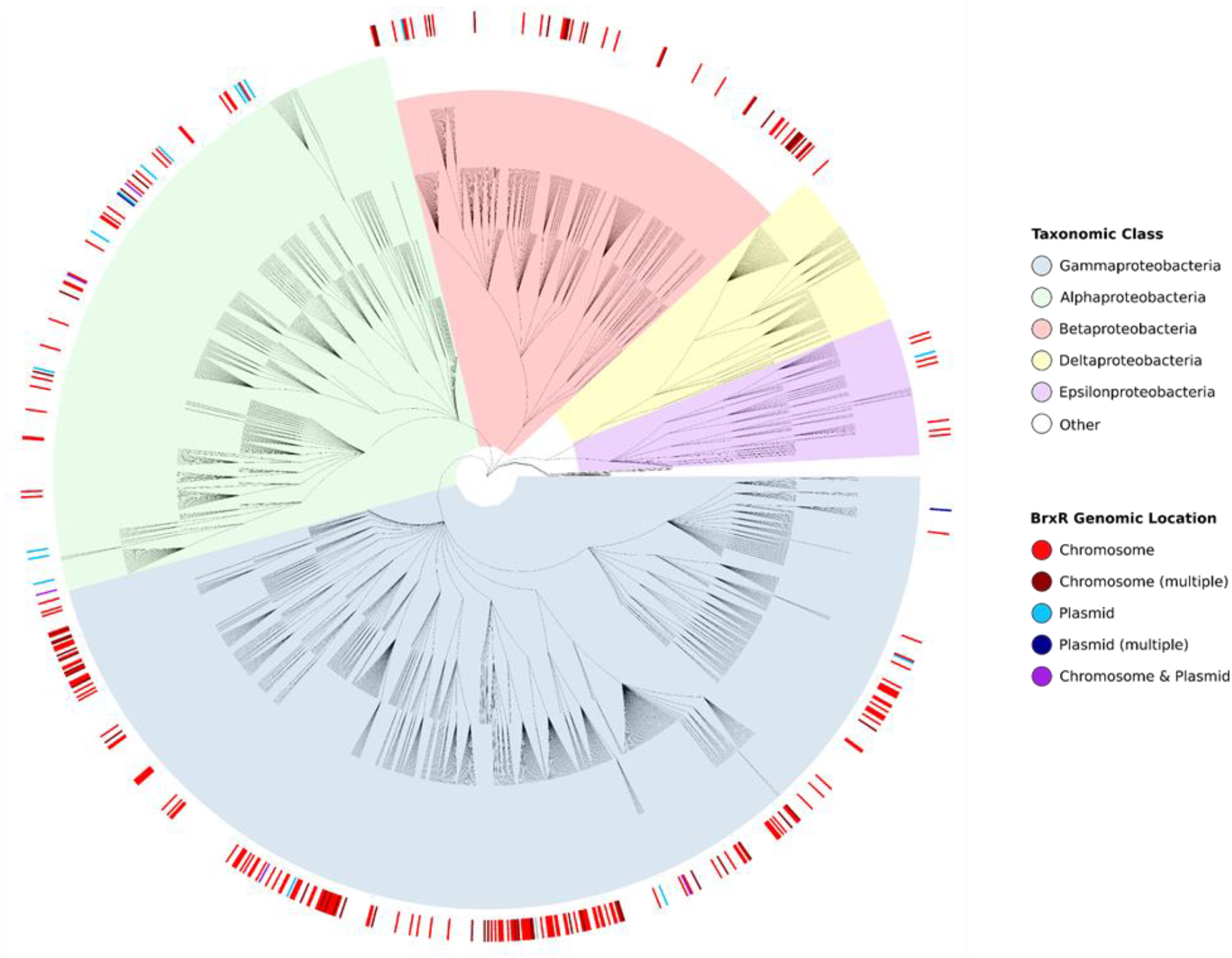
BrxR is widely distributed in the phylum proteobacter. Phylogenetic tree of 1589 proteobacterial genomes, as downloaded from the NCBI Taxonomy resource and background highlighted according to taxonomic class. BrxR hits are indicated in the exterior circle as a heatmap, coloured to show whether *brxR* is located on the chromosome or a plasmid.

### BrxR-family homologues are associated with diverse phage defence systems and islands in bacteria

Having identified a list of 347 BrxR homologues, we wanted to know how many phage defence systems could potentially be regulated. We compiled a list of 110 reference protein sequences, comprised of key genes from diverse known phage defence systems and sub-types (**Supplementary Table S3**), which was used to identify phage defence homologues within our database, using a BLASTP threshold of E-value < 1e-5. For each BrxR homologue, we tested for the presence of one or more phage defence homologues within 50 kb downstream of *brxR*, identifying 382 phage defence protein homologues. These 382 protein homologues were within 210 phage defence systems, and these were co-localised with 164 of 347 BrxR homologues (48.41%) (**Supplementary Table S4**). A less stringent threshold of E-value < 1e-3 was also tested, but this did not increase the number of BrxR homologues associated with known phage defence systems. We also examined the 50 kb upstream of each BrxR homologue, identifying a further 77/347 BrxR homologues including 29/347 with phage defence systems both upstream and downstream (**Supplementary Table S5**). This equates to an additional 48/347 BrxR homologues that were co-localised with at least one phage defence system, taking the total of associated BrxR homologues to 212/347 (61.10%). As BrxR_*Efer*_ controls phage defence systems downstream, we chose to be conservative and also focussed only on the downstream matches for further analysis.

Sorting the BrxR-associated phage defence systems by class showed BrxR homologues are predominantly co-localised with BREX systems (70/210 BrxR-associated phage defence systems, 33.33%; **Figure 6A**). Next, they are co-localised with type IV and type I restriction enzymes, 37/210 (17.61%) and 34/210 (16.19%), respectively. CRISPR-Cas systems were similarly well represented, comprising 21 of 210 BrxR-associated phage defence systems (10.00%). Whilst not all toxin-antitoxin system families have been shown to abort phage infections, there are multiple examples where toxin-antitoxin system types I-IV cause abortive infection (60–62). BrxR was predominantly co-localised with type II and IV toxin-antitoxin systems (**Figure 6A**). More recently defined phage defence systems such as Wadjet, Zorya, Thoeris (16), Pycsar (22) and CBASS (18) were also co-localised with BrxR homologues (**Figure 6A**). We visualised the BrxR-associated systems in relation to the host phylogeny (**Supplementary Figure S3**).

**Figure 6.**
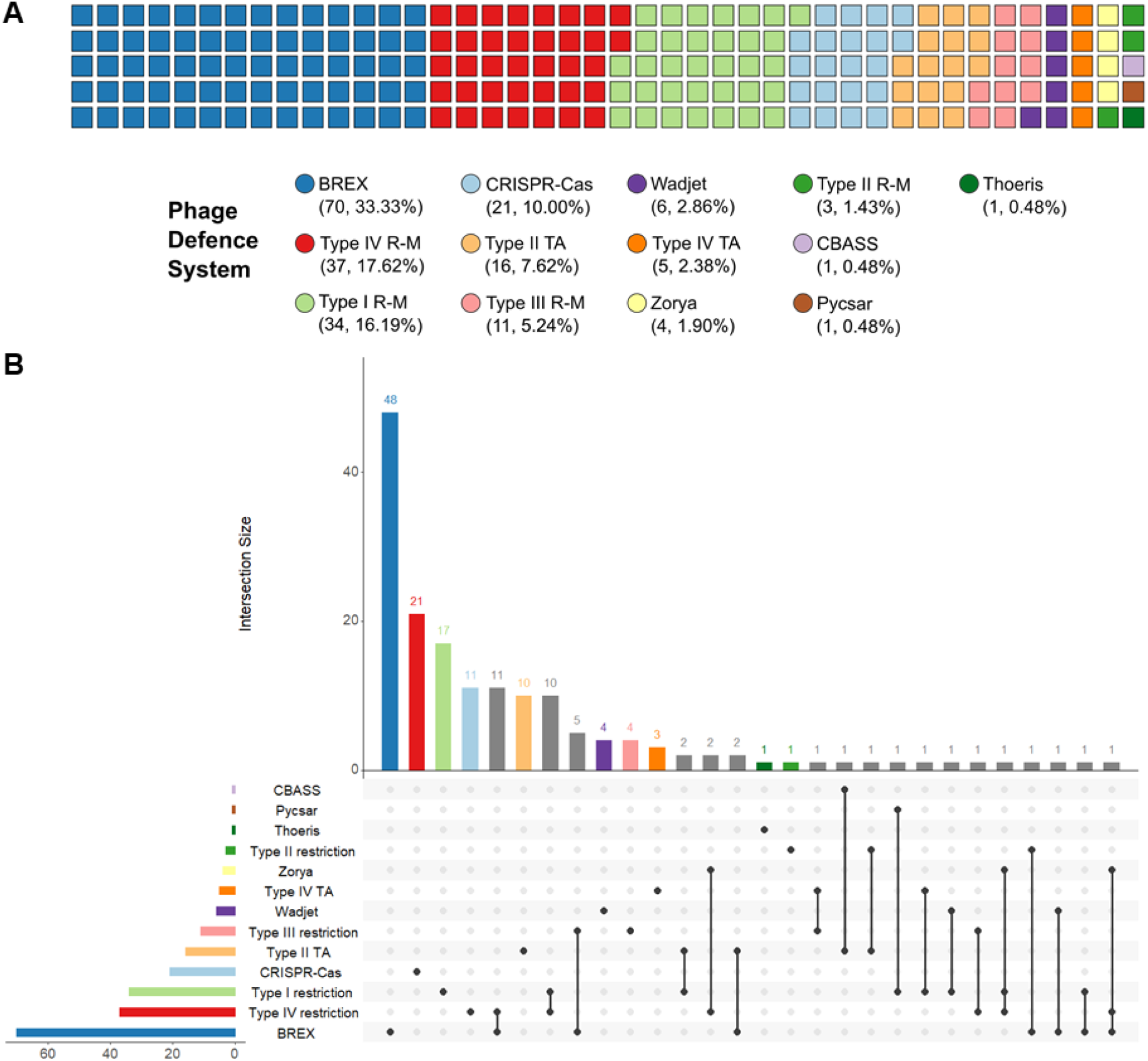
BrxR is associated with a variety of different phage defence systems. (**A**) Waffle chart showing the distribution of defence systems identified downstream of *brxR*, coloured to show the general class of system identified. (**B**) UpSet plot showing the co-occurrence of phage defence systems downstream of *brxR*. Set and intersection bars are coloured by single defence systems, with overlapping intersections in grey. The matrix indicates overlapping set intersections, with set size as the horizontal bar chart, and the vertical bar chart showing intersection size (the number of times any systems are found in combination).

Not only did our analysis identify individual phage defence systems that were associated with a BrxR homologue, but we also found examples of multiple systems, clustered into phage defence islands, that were associated with a BrxR homologue (**Figure 6B**). The most common clustering was between BREX and type IV restriction, as seen for pEFER and previously noted to be the most common pairing of phage defence systems (14, 15, 37). The next most common clusters included type III and type IV restriction systems, then BREX and type III restriction systems (**Figure 6B**). There were also individual examples of multiple forms of diverse clusters, including islands containing homologues from three different systems (**Figure 6B**). BrxR-associated phage defence systems were also further divided by sub-type of phage defence system (**Supplementary Figure S4**). Collectively, these data show how BrxR-homologues are likely used to regulate a wide range of phage defence islands that can be highly mosaic in phage defence system content, and within diverse hosts.

Our findings confirm that BrxR is the first member of a new family of transcriptional regulators involved in protecting bacteria from phages and mobile genetic elements.

## Discussion

Plasmid pEFER was known to encode a phage defence island encoding complementary BREX and type IV restriction systems (37). Here, we have shown that this defence island is regulated by a WYL-domain containing protein, BrxR_*Efer*_. BrxR_*Efer*_, together with the homologue BrxR_*Acin*_, investigated in a study co-submitted with this article (59), represent the first examples of a new family of transcriptional regulators.

BrxR_*Efer*_ acts as a transcriptional repressor, blocking transcription from a promoter upstream of *brxR* that controls the canonical first BREX gene, *brxA* (**Figure 1**). Whilst the BREX loci from *Bacillus cereus* contained another promoter upstream of *pglZ* (17), we were unable to detect a promoter in the comparable region of pEFER (**Figure 1**). EMSA studies demonstrated that BrxR_*Efer*_ bound as a stable dimer to inverted DNA repeats, in a sequence-dependent manner (**Figure 2**). These repeats were positioned immediately downstream of the predicted P_*brxR*_ promoter sequence, and so repression is likely due to sterically blocking the RNA polymerase. Curiously, there were no obvious IR sequences downstream of the predicted promoter sequences of P_*brxA*_, and so how this region is repressed by BrxR_*Efer*_ remains to be understood.

It is clear that not all BREX loci require a BrxR homologue (17, 63), so it is worth considering why transcriptional regulation of phage defence genes might be required. We suggest that some BREX homologues can be toxic, which could be exacerbated by the genomic context (chromosomal or plasmid-based) or the methylation status of the host. We have noted that PglX_*Efer*_ product is toxic when over-expressed and our repeated inability to make a *brxR*_Efer_ knockout mutant supports the hypothesis that the repression of BREX genes is required to reduce fitness costs to the host prior to phage infection.

BrxR proteins contain an N-terminal wHTH domain, a WYL-domain and a C-terminal dimerisation domain (**Figure 3**). The structures of BrxR_*Efer*_ and BrxR_*Acin*_ (59) are the first for this family, but increasing numbers of WYL-domain proteins have recently been characterised. The HTH-WYL protein Sll7009 has previously been shown to negatively regulate a CRISPR locus in *Synechocystis* (28). In this case, however, BrxR_*Efer*_ and Sll7009 share no significant sequence similarity. Further WYL-domain proteins hypothesised to be transcriptional regulators have also been identified through computational analyses of phage defence islands associated with integrative conjugative elements (34). WYL-domain containing proteins can also act as transcriptional regulators in contexts other than phage defence. For instance the HTH-WYL-WCX protein PafBC, which has a very different overall structure to BrxR_*Efer*_, is a transcriptional activator in response to DNA damage in mycobacteria (33). Similarly, the much larger DriD protein (914 amino acids to the 295 amino acids of BrxR_*Efer*_), contains HTH-WYL domains and is involved in upregulation of the DNA damage response in *C. crescentus* (29). WYL-domains can also play a role in regulating catalysis, with PIF1 helicase activity dependent on the WYL-domain (30), and Cas13d activity enhanced by the accessory WYL1 protein (31, 32).

It has previously been predicted that WYL domains could function as regulatory domains, either as switches to alter the activity of enzymes, or for transcriptional regulation, as part of phage defence (23). To perform such biological roles, the WYL-domains likely bind ligands; in PIF1, the WYL-domain binds ssDNA (30), whereas in WYL1 the domain binds ssRNA (32). To respond to DNA damage, it has been postulated that the WYL-domains bind ssRNA, ssDNA, or some other nucleic acid molecule or secondary messenger (29, 33). The reported promiscuity of WYL-domain ligand recognition will make it a challenge to identify the specific ligands experimentally. We speculate that a suitable candidate for recognising a phage infection might be a cyclic 2′-3′ phosphate, a cyclic nucleotide, or other nucleic acid polymers.

The structure of BrxR_*Efer*_ showed sulphate ions bound within the WYL-domain (**Figures 3**,**4**). This highly conserved fold is known to bind a large range of ligands (23, 24), and BrxR_*Efer*_ has an abundance of functional groups located in a conserved basic, solvent-exposed patch at the top of the WYL-domain, which are predicted to recognise the target ligand (**Figure 4**). We propose that ligand-binding alters the conformational state of BrxR to release the bound DNA, and de-repress transcription of phage defence genes. Interestingly, the structures of Efer_*Acin*_ (both *apo* and DNA-bound), present a C-terminal strap extending back over the protomeric WYL-domain, perhaps indicating some form of lid mechanism that regulates ligand recognition and binding (59). It is clear that future systematic analysis of potential ligands, combined with extensive mutagenesis studies, are required to identify the molecules that bind BrxR, and determine whether they do cause de-repression.

Comparative genomic analyses identified a larger family of BrxR homologues, widespread within Proteobacteria (**Figure 5**). Stringent thresholds were used to exclude the many prokaryotic WYL-domain containing proteins. Attempts to match these BrxR-family homologues with known phage defence systems demonstrated that nearly half were associated with a diverse array of single defence systems, or a variety of collections of systems within defence islands (**Figure 6**). BREX systems and type IV restriction enzymes were most highly represented, consistent with previous studies that show this pairing was the most prevalent in defence islands (14, 15).

In addition, BrxR-homologues were associated with a large array of other systems and islands, suggesting that BrxR-homologues might not simply function to avoid fitness costs (as hypothesised above), but also to regulate the time-course or stages of phage defence. Such a mechanism would allow each system to provide protection, depending on context of infection and the counter-defence systems present on the invading phage. The possibility of phage-dependent ligands binding to BrxR also provides an opportunity for phage defence systems or islands to respond dynamically to the type of attack, be it via a phage, or a mobile genetic element. In this manner, only selected invading DNAs (or RNAs) might be targeted.

Because almost half of the BrxR-family homologues were associated with known phage defence systems, there is an exciting possibility that other conserved genes associated with BrxR-family homologues represent core genes of yet undiscovered phage defence systems. By using BrxR to hunt for new systems, it may be possible to further expand our knowledge of phage-host interactions and identify novel tools for biotechnology.

## Data Availability

The crystal structure of BrxR has been deposited in the Protein Data Bank under accession number 7QFZ. All other data needed to evaluate the conclusions in the paper are present in the paper and/or Supplementary Data.

## Acknowledgements

We gratefully acknowledge Diamond Light Source for time on beamlines I24 and I04 under proposal MX24948. We thank David Dryden for his invaluable input on the project and manuscript. We thank Nicolas Wenner for training.

## Supplementary Figures

**Supplementary Figure S1.**
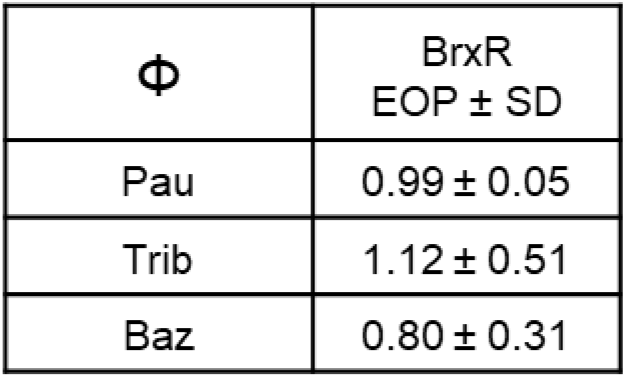
BrxR does not provide phage defence. EOP values for two BREX-sensitive phages (Pau, Trib) and one BREX-resistant phage (Baz), were determined against DH5α pBAD3O-*hiS*_*6*_-*brxR*, with the induced plasmid as test strain, and uninduced plasmid as control. Values shown are mean and standard deviation of triplicate data.

**Supplementary Figure S2.**
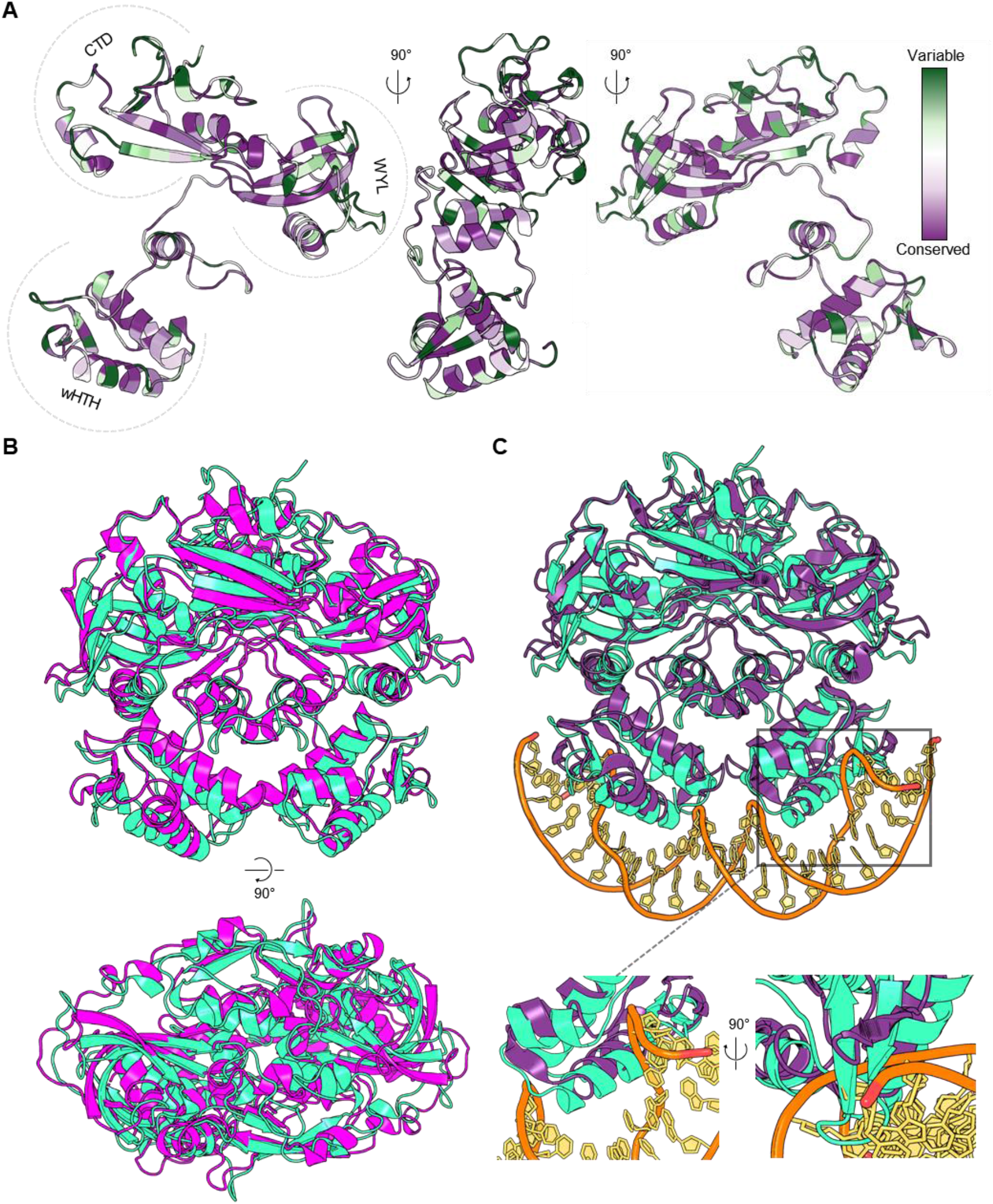
Conserved residues and structures within the BrxR-family. **(A)** Consurf analysis of BrxR_*Efer*_ shows residue conservation over a single protomer. Orthogonal views are shown and are indicated by directional arrows. Conservation is indicated by a colour gradient from highly variable (green) to highly conserved (purple). **(B)** An alignment of the apo structures of BrxR_*Efer*_ (cyan) and BrxR_*Acin*_ (magenta) (PDB 7T8L) shows the same overall fold with variations in secondary structure positioning. Orthogonal views are shown as indicated by the directional arrow. **(C)** An alignment of *apo* BrxR_*Efer*_ (cyan) and DNA-bound BrxRA_*Acin*_(purple) (PDB 7T8K) clearly shows differences within the wHTH domains. A close-up orthogonal views of the solid boxed region demonstrates the clashing of the wing of BrxR_*Efer*_with DNA.

**Supplementary Figure S3.**
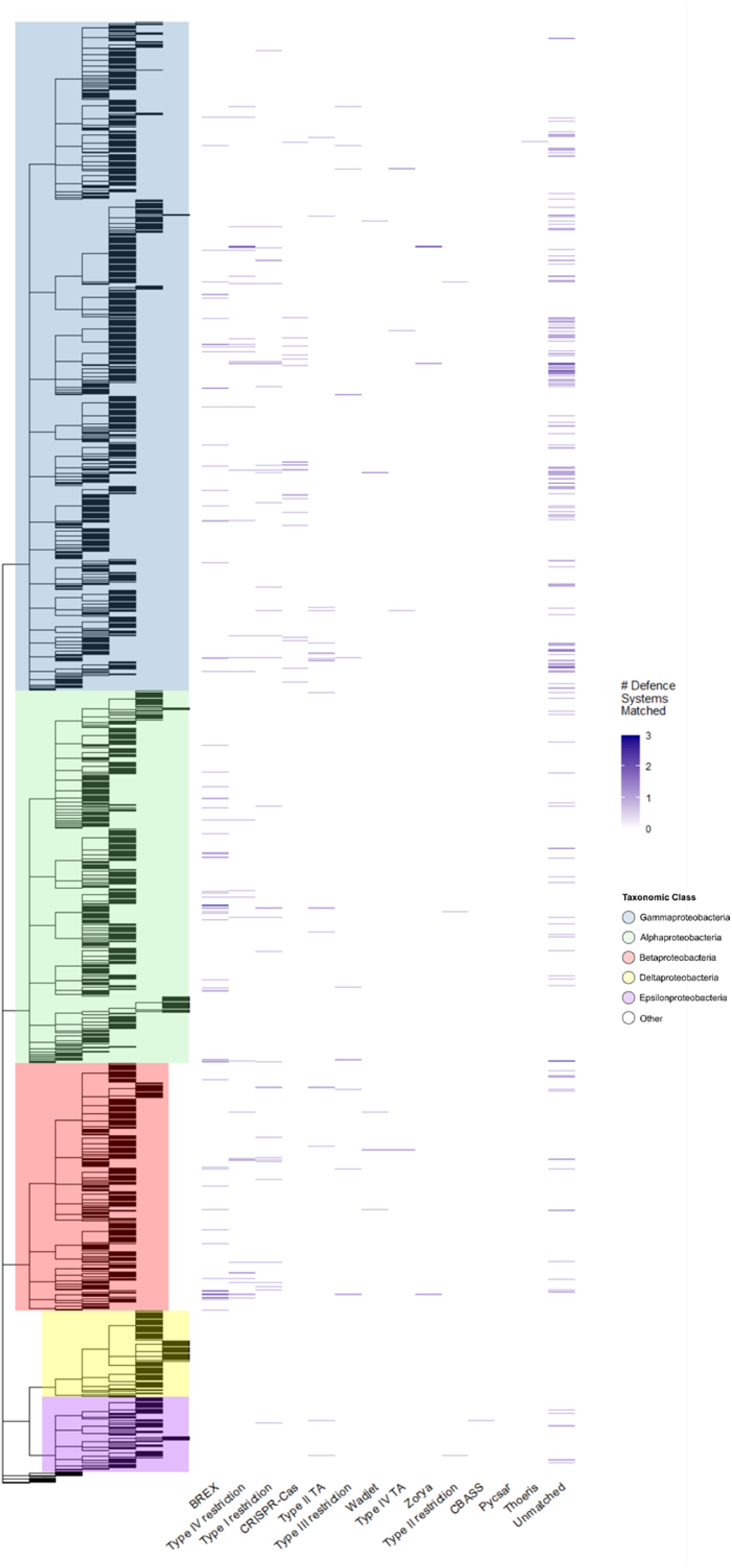
Heatmap of BrxR hits showing the distribution of associated phage defence systems within proteobacteria. Represented phage defence systems are distributed according to host phylogeny. Scale indicates number of matched defence systems per species. The phylogenetic distribution of 51.59% of BrxR-family homologues not associated with known phage defence systems are shown in the column labelled Unmatched.

**Supplementary Figure S4.**
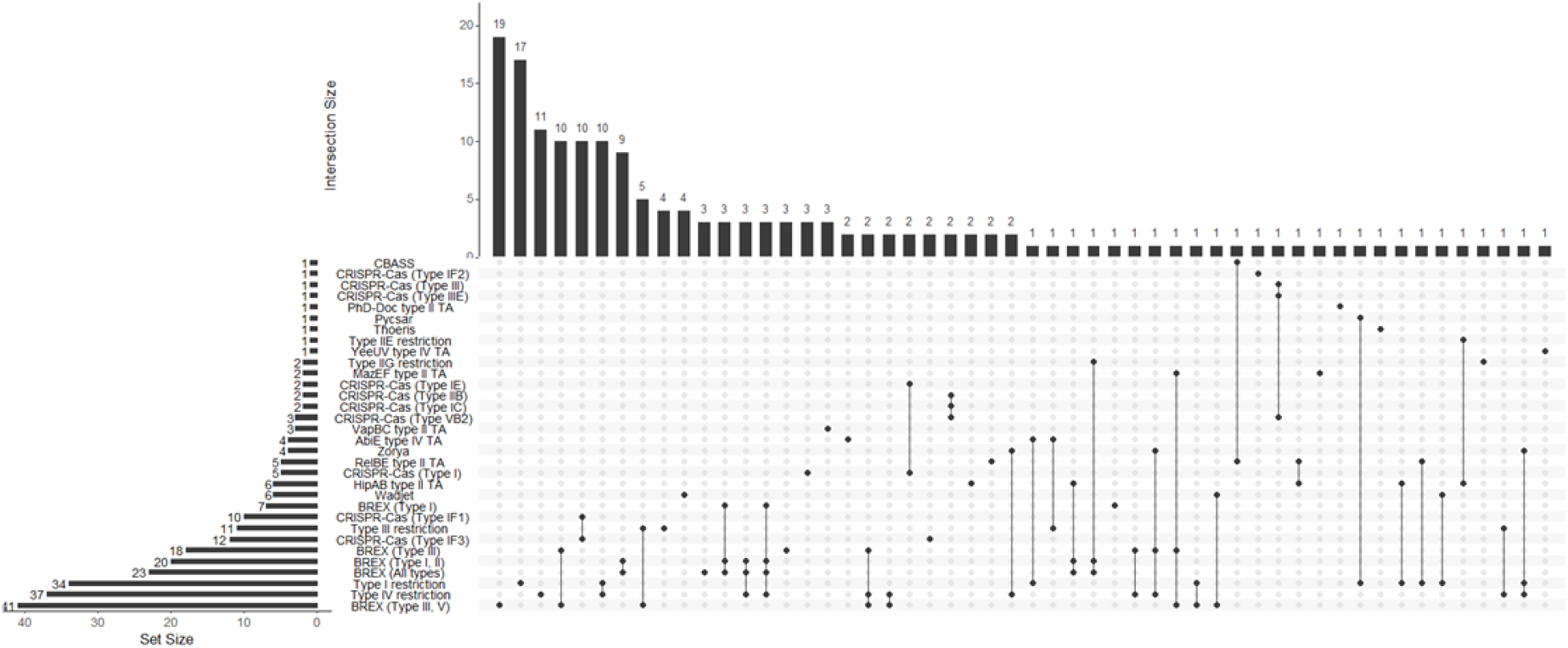
Extended UpSet plot of specific phage defence systems identified downstream of *brxR*. The matrix indicates overlapping set intersections, with set size as the horizontal bar chart, and the vertical bar chart showing intersection size (the number of times any systems are found in combination).

## Supplementary Tables

**Supplementary Table S1.**
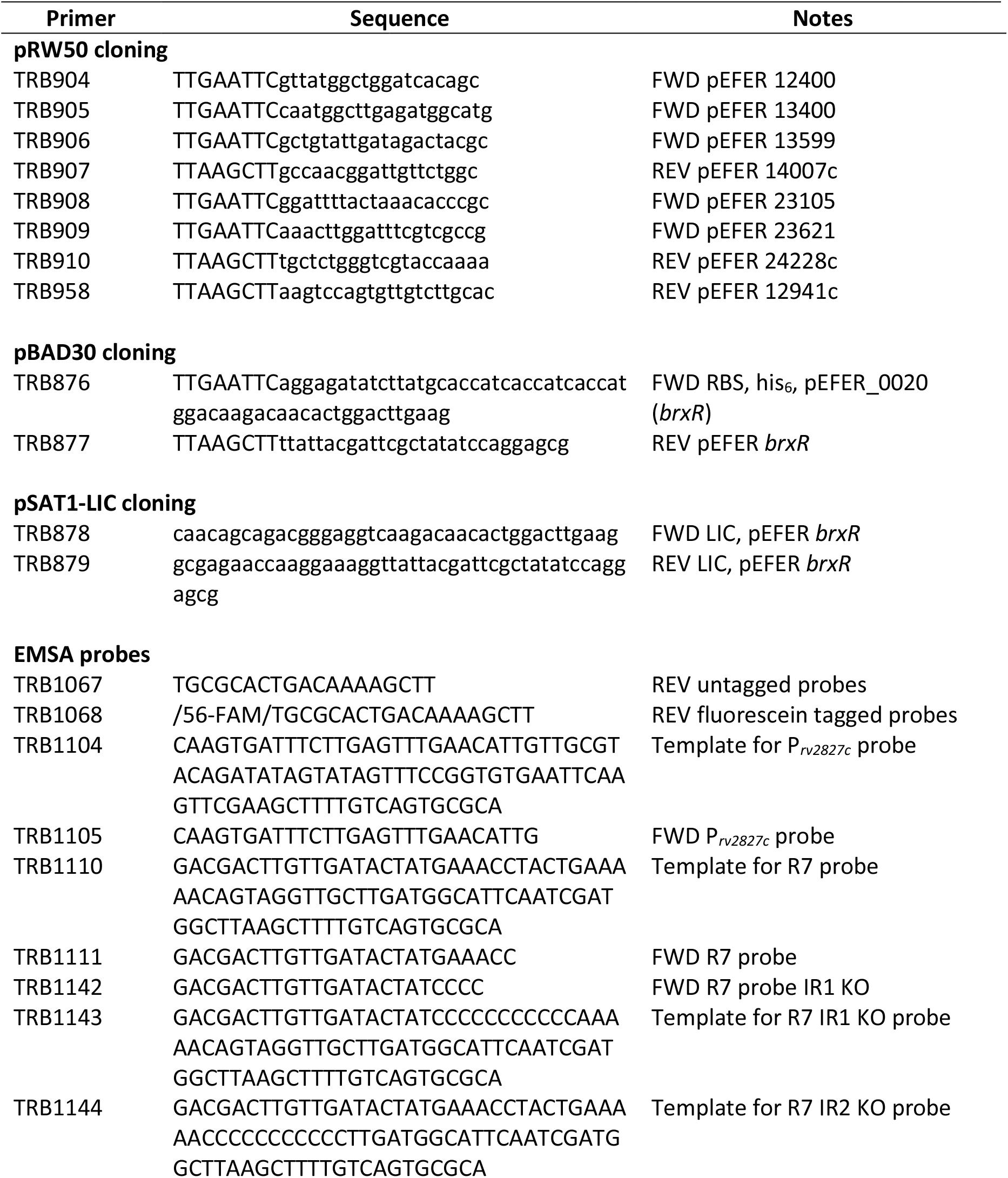
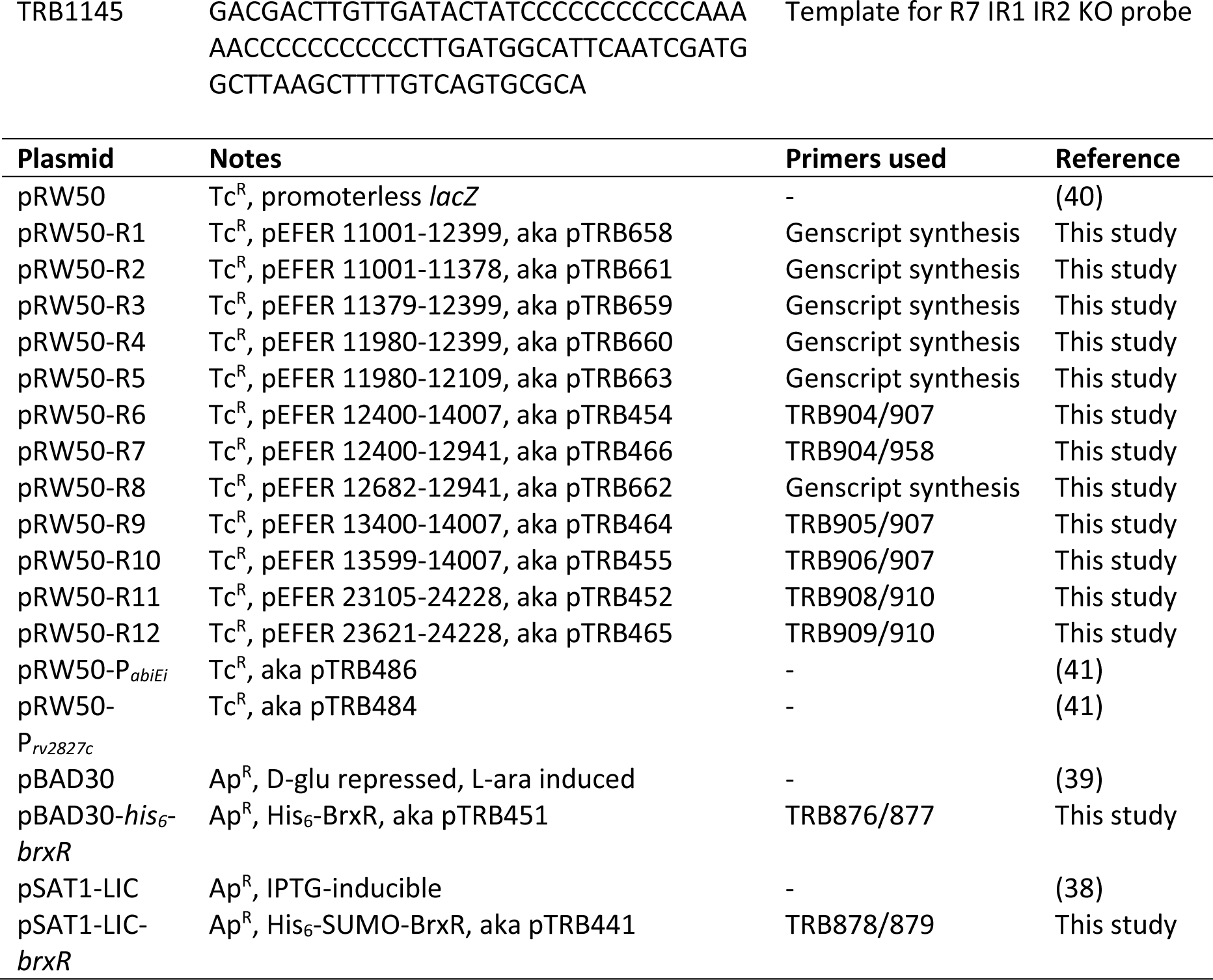
Oligonucleotides and plasmids used in this study.

**Supplementary Table S2.** List of DALI hits (excel)

**Supplementary Table S3.** Reference phage defence protein sequences.

| **Protein** | **Associated with Phage-Defence System** | **Accession** | **Reference** |
| --- | --- | --- | --- |
| BrxR | NA | WP_017044131.1 | Picton et al., 2021 |
| PglZ | BREX (All types) | WP_004722530.1 | Goldfarb et al., 2015 |
| BrxC | BREX (All types) |  | Goldfarb et al., 2015 |
| Cas3 | CRISPR-Cas (Type I) | WP_037623090.1 | Makarova et al., 2019 |
| Cas9 | CRISPR-Cas (Type II) | WP_024703962.1 | Makarova et al., 2019 |
| Cas10 | CRISPR-Cas (Type III) | WP_014621547.1 | Makarova et al., 2019 |
| PglW | BREX (Type II) | NP_630703.1 -<br>WP_011031052.1 | Goldfarb et al., 2015 |
| BrxP | BREX (Type IV) | YP_001716949.1 -<br>WP_012301903.1 | Goldfarb et al., 2015 |
| BrxHII | BREX (Type III, V) | YP_004121416.1 -<br>WP_013514587.1 | Goldfarb et al., 2015 |
| PglX | BREX (Type I, II) | WP_001095615.1 | Goldfarb et al., 2015 |
| PglXI | BREX (Type III) | YP_004121414.1 –<br>WP_013514585.1 | Goldfarb et al., 2015 |
| BrxU (GmrSD) | Type IV restriction | WP_000283751.1 | Picton et al., 2021 |
| SspC | SspABCD-SspE | WP_016789109.1 | Xiong et al., 2020 |
| DndD | DndABCDE-FGH | WP_042527205.1 | Wang et al., 2007 |
| PvuRts1I | Type IV restriction | WP_011039664.1 | Kazrani et al., 2014 |
| Mrr | Type IV restriction | WP_000217936.1 | Loenen and Raleigh, 2014 |
| McrA | Type IV restriction | WP_000557907.1 | Loenen and Raleigh, 2014 |
| McrB | Type IV restriction | WP_000379041.1 | Loenen and Raleigh, 2014 |
| SauUSI | Type IV restriction | WP_038539189.1 | Loenen and Raleigh, 2014 |
| ScoA3I | Type IV restriction | WP_011030182.1 | Loenen and Raleigh, 2014 |
| MspJI | Type IIM restriction | WP_188871137.1 | Loenen and Raleigh, 2014 |
| DpnI | Type IIM restriction | WP_000418960.1 | Loenen and Raleigh, 2014 |
| MmeI | Type IIL restriction | WP_018986935.1 | Callahan et al., 2016 |
| EcoKI | Type I restriction | WP_001272447.1 | Loenen et al., 2014 |
| EcoprrI | Type I restriction | CAA36526.1 | Loenen et al., 2014 |
| KpnBI | Type I restriction | AAA97402.1 | Loenen et al., 2014 |
| M.StySKI | Type I restriction | EIQ50780.1 | Loenen et al., 2014 |
| Eco57I | Type IIG restriction | WP_032180232.1 | Loenen et al., 2014 |
| Bcgl | Type IIB restriction | WP_013853237.1 | Loenen et al., 2014 |
| SapI | Type IIA restriction | TSC79897.1 | Pingoud et al., 2014 |
| BpuSI | Type IIC restriction | WP_098381443.1 | Pingoud et al., 2014 |
| EcoRII | Type IIE restriction | WP_001532073.1 | Pingoud et al., 2014 |
| Cfr10I/Bse634I | Type IIF restriction | WP_072444636.1 | Pingoud et al., 2014 |
| DptF | Type IIH /<br>phosphorothioation-<br>dependent restriction | WP_000417613.1 | Xu et al., 2020 |
| AhdI | Type IIH restriction | WP_168235528.1 | Pingoud et al., 2014 |
| EcoRI | Type IIP restriction | WP_001565219.1 | Pingoud et al., 2014 |
| FokI | Type IIS restriction | AAA24927.1 | Pingoud et al., 2014 |
| BbvCI | Type IIT restriction | AAX14652.1 | Pingoud et al., 2014 |
| EcoP15I | Type III restriction | WP_032190829.1 | Rao et al., 2014 |
| LlaFI | Type III restriction | AAD15793.1 | Rao et al., 2014 |
| PstII | Type III restriction | AAZ73167.1 | Rao et al., 2014 |
| StyLT1 | Type III restriction | AAB26534.1 | Rao et al., 2014 |
| DUF4435 | PARIS | WP_001696664.1 | Rousset et al., 2020 |
| BstA | BstA | WP_000248006.1 | Owen et al., 2021 |
| ZorA | Zorya | WP_186701292.1 | Doron et al., 2018 |
| DrmA | DISARM | WP_227539820.1 | Ofir et al., 2017 |
| pVip8 | Viperins | WP_019672856.1 | Bernheim et al., 2021 |
| NucC | CBASS | WP_001286625.1 | Lau et al., 2020 |
| Cyclase | Pycsar | WP_053265929.1 | Tal et al., 2021 |
| DncV | cGAS | WP_001901330.1 | Cohen et al., 2019 |
| Eco8 | Retron-Eco8 | WP_023304001.1 | Millman et al., 2020 |
| AbiEii | AbiE type IV TA | WP_041980506.1 | Dy et al., 2014 |
| ToxN | ToxIN type III TA | WP_012609144.1 | Fineran et al., 2009 |
| MazF | MazEF type II TA | WP_000254738.1 | Hazan and Engelberg-Kulka, 2004 |
| Hok | Hok/Sok type I TA | WP_001302699.1 | Pecota and Wood, 1996 |
| ThsB | Thoeris | WP_071563695.1 | Doron et al., 2018 |
| JetC | Wadjet | WP_176335405.1 | Doron et al., 2018 |
| Csa5_IA | CRISPR-Cas (Type IA) | WP_010879363.1 | Makarova et al., 2019 |
| Cas8b1_IB | CRISPR-Cas (Type IB) | WP_012103098.1 | Makarova et al., 2019 |
| Cas8c_IC | CRISPR-Cas (Type IC) | WP_010896519.1 | Makarova et al., 2019 |
| Csc3_ID | CRISPR-Cas (Type ID) | ACK70148.1 | Makarova et al., 2019 |
| Cse1_IE | CRISPR-Cas (Type IE) | WP_046891358.1 | Makarova et al., 2019 |
| Csy1_IF1 | CRISPR-Cas (Type IF1) | WP_050296085.1 | Makarova et al., 2019 |
| Cas7f2_IF2 | CRISPR-Cas (Type IF2) | WP_011919225.1 | Makarova et al., 2019 |
| Csy3_IF3 | CRISPR-Cas (Type IF3) | WP_048664206.1 | Makarova et al., 2019 |
| Csb2_IU | CRISPR-Cas (Type IU/IG) | WP_010940732.1 | Makarova et al., 2019 |
| Csf1_IVA | CRISPR-Cas (Type IVA) | ADC73187.1 | Makarova et al., 2019 |
| Csf2_IVB | CRISPR-Cas (Type IVB) | WP_029308491.1 | Makarova et al., 2019 |
| Cas5_IVC | CRISPR-Cas (Type IVC) | RME36479.1 | Makarova et al., 2019 |
| Csm2_IIIA | CRISPR-Cas (Type IIIA) | AAW53329.1 | Makarova et al., 2019 |
| Csm1_IIIA | CRISPR-Cas (Type IIIA) | WP_002486045.1 | Makarova et al., 2019 |
| Cmr1_IIIB | CRISPR-Cas (Type IIIB) | WP_011012270.1 | Makarova et al., 2019 |
| Cmr2_IIIB | CRISPR-Cas (Type IIIB) | WP_011012269.1 | Makarova et al., 2019 |
| Cmr3_IIIB | CRISPR-Cas (Type IIIB) | WP_011012268.1 | Makarova et al., 2019 |
| Cmr4_IIIC | CRISPR-Cas (Type IIIC) | WP_012003242.1 | Makarova et al., 2019 |
| Cas10_IIID | CRISPR-Cas (Type IIID) | WP_011153736.1 | Makarova et al., 2019 |
| Csx10_IIID | CRISPR-Cas (Type IIID) | WP_099887203.1 | Makarova et al., 2019 |
| Cas7_IIIE | CRISPR-Cas (Type IIIE) | KHE91659.1 | Makarova et al., 2019 |
| Csm3_IIIF | CRISPR-Cas (Type IIIF) | WP_012002152.1 | Makarova et al., 2019 |
| Cas9_IIA | CRISPR-Cas (Type IIA) | WP_011227028.1 | Makarova et al., 2019 |
| Cas1_IIB | CRISPR-Cas (Type IIB) | WP_011212793.1 | Makarova et al., 2019 |
| Cas2_IIC1 | CRISPR-Cas (Type IIC1) | WP_002214566.1 | Makarova et al., 2019 |
| Cas2_IIC2 | CRISPR-Cas (Type IIC2) | OJI07265.1 | Makarova et al., 2019 |
| Cpf1_VA | CRISPR-Cas (Type VA) | WP_014550095.1 | Makarova et al., 2019 |
| Cas12b_VB1 | CRISPR-Cas (Type VB1) | WP_021296342.1 | Makarova et al., 2019 |
| Cas4_VB2 | CRISPR-Cas (Type VB2) | QBM02857.1 | Makarova et al., 2019 |
| Cas12c_VC | CRISPR-Cas (Type VC) | KZX85786.1 | Makarova et al., 2019 |
| Cas12d_VD | CRISPR-Cas (Type VD) | OJI08769.1 | Makarova et al., 2019 |
| Cas14a_VF1 | CRISPR-Cas (Type VF1) | QBM01136.1 | Makarova et al., 2019 |
| Cas13a_VIA | CRISPR-Cas (Type VIA) | WP_018451595.1 | Makarova et al., 2019 |
| Cas13d VID | CRISPR-Cas (Type VID) | WP_215648980.1 | Makarova et al., 2019 |
| Csx28_VIB1 | CRISPR-Cas (Type VIB1) | WP_115154074.1 | Makarova et al., 2019 |
| PglZ_B | BREX Type I | WP_089368483.1 | Goldfarb et al., 2015 |
| EndoR | Type I restriction | WP_153250578.1 | NA |
| VapC | VapBC type II TA | WP_153250579.1 | NA |
| EndoS | Type I restriction | WP_004745828.1 | NA |
| YeeU | YeeUV type IV TA | WP_012154510.1 | Masuda et al., 2012 |
| AbiEii_2 | AbiE type IV TA | WP_012368380.1 | Dy et al., 2014 |
| RelE | RelBE type II TA | WP_014703508.1 | NA |
| EndoM | Type I restriction | WP_015879339.1 | NA |
| VapC_2 | VapBC type II TA | WP_043869448.1 | NA |
| HipA | HipAB type II TA | WP_061904595.1 | NA |
| AbiEi | AbiE type IV TA | WP_081921101.1 | Dy et al., 2014 |
| BrxU_2 | Type IV restriction | WP_089067936.1 | Picton et al., 2021 |
| RelE_2 | RelBE type II TA | WP_095699740.1 | NA |
| BrxF | BREX (Type III) | WP_107219702.1 | Goldfarb et al., 2015 |
| PhD | PhD-Doc type II TA | WP_128384231.1 | NA |
| AbiEii_3 | AbiE type IV TA | WP_142820193.1 | Dy et al., 2014 |
| BrxU_3 | Type IV restriction | WP_178969665.1 | Picton et al., 2021 |
| EndoR_2 | Type I restriction | WP_191600125.1 | NA |
| Cas2_IC | CRISPR-Cas (Type IC) | WP_199275886.1 | Nam et al., 2012 |

**Supplementary Table S4**. List of BrxR hits and associated phage defence systems (excel)

**Supplementary Table S5**. List of additional upstream BrxR-associated phage defence systems (excel)

